# Survey, Identification and Evaluation of biodiversity of *Seabuckthorn* (*Hippophae salicifolia*) in hills of Uttarakhand

**DOI:** 10.1101/2020.10.26.354951

**Authors:** Ranjit Singh, S.K. Dwivedi, Madhu Bala

**Affiliations:** Defence Institute of Bio-Energy Research (DRDO), Haldwani, Uttarakhand

**Keywords:** Biodiversity, Uttarakhand, Seabuckthorn, Morphotypes, Field gene bank

## Abstract

Extensive studies were carried out during 2017-2019 to evaluate the *Hippophae salicifolia* (Seabuckthorn) biodiversity in Uttarakhand for their physico-chemical/biochemical parameters of ripe fruits like acidity (1.78-3.01 %), pH (2.15-2.87), Vitamin ‘C’ (271-494mg/100g), Total soluble solids (5.1-9.8) etc. Variation was also observed in fruit color (light yellow, yellow, light yellow orange, orange and orange red), fruit size (length 6.29-8.77mm, width 4.80-7.59mm), Juice (44.23-76.18%), Pomace (8.01-28.30%) and seed (6.8-21.1%). On the basis of above parameters, promising morphotypes were identified and conserved at field gene bank at DIBER field station, Auli (Joshimath) 3142m AMSL.

## Introduction

*Hippophae salicifolia* commonly known as Seabuckthorn is one of the important wild plants of Uttarakhand Himalayas with nutritious fruits which are having tremendous medicinal & pharmaceutical importance. Besides Uttarakhand it is also found occurring naturally in other parts of India including Himachal Pradesh, Sikkim and Arunachal Pradesh. It is locally known as ‘*Ames*’, ‘*Ameel*’ and ‘*Chuk*’. *Hippophae salicifolia* belongs to family *Eleagnaceae* and is commonly known as willow leaved sea buckthorn and its fruits are very rich in vitamins and other important bio-chemicals or biologically active compounds (Rongsen, 1992).

In India, three species of *Hippophae* are reported, *H. rhamnoides* and *H. salicifolia* are common but third one *Hippophae tibetana* is found growing in very limited patches in high altitude hills (Singh et al., 2005). Among common two species, *H. salicifolia* has not been studied much for its high value medicinal benefits (Gupta et al., 2011). It differs from *H. rhamnoides* in two major aspects. First one is that it is a shrub that could grow up to a tree size (4–10m) at 1500–3200m above mean sea level and is limited in its bio-geographical distribution to the Himalayas (central and northern) (Gupta et al., 2011; Kaushal et al., 2013). *H. rhamnoides*, on the other hand is a bushy tree that is distributed widely in both Asia and Europe at higher altitudes (R. Singh, 2004; Chakraborty et al., 2016; Usha et al., 2014). It is found in Tibet region of China, Bhutan, Nepal, and Himalayan hills of India (Ramu et al., 2014). In India, it dwells at high altitude regions of Himachal Pradesh, Jammu and Kashmir, Sikkim, and Uttarakhand (Usha et al., 2014). Indigenously, the ethno-botanical uses of *H. salicifolia* by the regional people of Central Himalaya include animal feed, cosmetics, food, fuel, medicine, veterinary care, and bio-fencing (Singh et.al, 2005; Thakur et al., 2015).

In Uttarakhand this plant is found abundantly in higher hills of Chamoli and Uttarkashi Districts, but not much biodiversity study has been done so far, hence a systematic study in Chamoli District was carried out by the Defence Institute of Bio-energy Research, Haldwani to study the existing biodiversity in the region.

## Materials & Methods

The study was undertaken by Defence Institute of Bio-Energy Research, Haldwani, having a field station at Auli (11000 ft) above mean sea level where Seabuckthorn R & D is being carried out. During the extensive survey Random as well as biased sampling was followed (Sinha 1981) to collect particular morphotypes. The physico-chemical properties of ripe fruits were analyzed within 48 hours of collection. The fruit colour was analyzed as per Royal Horticultural Society colour chart (RHS 2001). Fruit size i.e. length and width were measured by digital vernier calipers while fruit weight was measured by digital weighing balance. The volume was calculated by using water replacement method. Total soluble solids (TSS) was measured with digital refractometer and pH of the juice was recorded by digital pH meter. Other parameters like acidity (as malic acid), ascorbic acid and sugar content of the fruit juice were estimated as per Rangana (1986), for calculations standard statistical procedures were followed. Entire data was pooled and analyzed as per standard statistical procedures (Panse and Sukhatme 1986)

## Results & Discussion

DIBER, Haldwani has conducted the first ever extensive systematic survey in the hills of Uttarakhand. The natural habitat of Seabuckthorn identified are Yamunotri, Gomukh, Harhindum, Hanuman chatti, Badrinath area, Urgam valley, Jelam area of Malari, at an altitudinal range of 2700-3700m above msl in Uttarakhand hills (Figure 1). *H. salicifolia* is generally found growing naturally along the rivers, channels, tributaries and along roadside in some places. It has been found growing in hill slopes, rocky and sandy soils. Major Seabuckthorn growing areas were found in Mana, Badrinath Hanuman Chatti and Bhunder along the Alaknanda and its tributaries. Its habit is generally a tree with an average height of 2-12m but sometimes bushy habit plants were also observed. Most of the seabuckthorn population is natural one and thick habitat is due to profuse suckering ability of the plant and its suckers run parallelly and give rise to new plants at intervals. Seabuckthorn being a cross pollinated plant lot of variation in the plant population has been noticed in the region.The data pertaining to Physico-chemical properties of mature fruits of *Hippophae salicifolia* collected from various locations have been presented in Table-1-6.

**Table-1:** Physico-chemical characteristics of fruits of *Hippophae salicifolia* morphotypes from Badrinath region in Uttarakhand

| S.No | Sample | Altitude (m) | Fruit colour And code | Fruit Shape | Fruit length (cm) | Fruit width (cm) | TSS (°B) | Juice (%) | Seed (%) | Pomace (%) | 100 fruit weight (g) | 100 fruit Volume (ml) | pH | Vitamin 'C' mg/100 g | Acidity % mg/100 g | Total Sugar mg/100 g |
| --- | --- | --- | --- | --- | --- | --- | --- | --- | --- | --- | --- | --- | --- | --- | --- | --- |
| 1 | DIBER-1 | 3137 | YG | Round | 6.56 | 6.48 | 6.0 | 59.67 | 9.65 | 30.68 | 29.3 | 14.0 | 2.4 | 352 | 2.19 | 2.87 |
| 2 | DIBER-2 | 3137 | YG | Oval | 6.33 | 6.32 | 5.3 | 54.34 | 10.1 | 35.56 | 22.0 | 11.0 | 2.1 | 325 | 2.28 | 3.11 |
| 3 | DIBER-3 | 3137 | LY | Capsular | 7.00 | 6.50 | 6.1 | 59.11 | 9.45 | 31.44 | 35.9 | 17.5 | 2.3 | 423 | 2.69 | 3.18 |
| 4 | DIBER-4 | 3137 | LG | Elliptical | 7.98 | 6.65 | 5.6 | 64.42 | 7.90 | 27.68 | 39.3 | 19.0 | 2.8 | 364 | 2.41 | 2.98 |
| 5 | DIBER-5 | 3137 | LG | Round | 7.55 | 5.31 | 6.3 | 61.21 | 9.12 | 29.67 | 32.8 | 15.6 | 2.7 | 372 | 2.33 | 2.86 |
| 6 | DIBER-6 | 3046 | LY | Round | 7.48 | 6.13 | 6.1 | 77.17 | 14.8 | **8.01** | 28.5 | 14.0 | 2.5 | 312 | 2.52 | 3.11 |
| 7 | DIBER-7 | 3046 | YO | Oval | 8.75 | 6.11 | 6.4 | 69.18 | 20.4 | 10.42 | 43.1 | 20.2 | 2.6 | 396 | 2.78 | 3.21 |
| 8 | DIBER-8 | 3046 | LG | Oval | 6.29 | 4.80 | 7.2 | 68.24 | 18.3 | 13.46 | **17.5** | 10.0 | 2.4 | 474 | 1.98 | 3.08 |
| 9 | DIBER-9 | 3046 | LY | Round | 6.52 | 6.31 | 7.2 | 71.54 | **6.08** | 22.38 | 25.9 | 13.0 | 2.3 | 301 | 2.28 | 3.09 |
| 10 | DIBER-10 | 3046 | LY | Oval | 6.29 | 4.80 | 7.2 | **44.21** | 20.9 | 34.89 | 28.8 | 13.5 | 2.4 | 376 | 2.21 | 2.99 |
| 11 | DIBER-11 | 3046 | YG | Round | 7.15 | 5.43 | 7.4 | 55.32 | 18.2 | 26.48 | 27.2 | 12.8 | 2.1 | 403 | 1.92 | 2.85 |
| 12 | DIBER-12 | 3046 | LG | Oval | 6.90 | 5.99 | 6.1 | 56.52 | 19.6 | 23.88 | 32.4 | 14.2 | 2.3 | 356 | 2.05 | 3.25 |
| 13 | DIBER-13 | 2976 | LY | Round | 7.05 | 6.12 | 6.4 | 53.26 | 20.4 | 26.34 | 34.8 | 16.8 | 2.8 | 322 | 2.27 | 3.69 |
| 14 | DIBER-14 | 2976 | LY | Oval | 6.89 | 6.08 | 6.2 | 58.24 | 14.8 | 26.96 | 24.5 | 12.0 | 2.7 | 437 | 1.95 | 3.56 |
| 15 | DIBER-15 | 2976 | LY | Round | 6.46 | 6.34 | 5.6 | 54.23 | 18.6 | 27.17 | 25.8 | 10.0 | 2.5 | 376 | 3.01 | 3.21 |
| 16 | DIBER-16 | 2976 | LY | Round | 6.13 | 5.93 | 6.1 | 49.71 | 19.6 | 30.69 | 27.3 | 13.0 | 2.6 | 305 | 2.15 | **2.73** |
| 17 | DIBER-17 | 2976 | LG | Oval | 6.45 | 6.14 | 5.6 | 51.45 | 18.4 | 30.15 | 28.9 | 13.2 | 2.4 | 437 | 2.33 | 3.01 |
| 18 | DIBER-18 | 2976 | LY | Round | 8.13 | 6.12 | 8.3 | 50.57 | 16.8 | 32.63 | 24.1 | 10.0 | 2.3 | 444 | 2.82 | 3.23 |
| 19 | DIBER-19 | 2976 | LG | Round | 7.37 | 6.32 | 6.6 | 50.56 | 16.4 | 33.04 | 30.2 | 12.8 | 2.4 | 437 | 2.36 | 3.67 |
| 20 | DIBER-20 | 2874 | LY | Round | 7.87 | 5.34 | 6.5 | 51.25 | 17.9 | 30.85 | 27.1 | **10.0** | 2.1 | 464 | 2.95 | 3.11 |
| 21 | DIBER-21 | 2874 | LG | Round | 7.24 | 5.69 | 7.1 | 54.21 | 18.2 | 27.59 | 28.3 | 10.2 | 2.3 | 366 | 2.08 | 2.89 |
| 22 | DIBER-22 | 2874 | LY | Oval | 7.14 | 6.02 | 8.3 | 56.12 | 19.6 | 24.28 | 30.7 | 11.0 | 2.4 | 362 | 2.30 | 2.97 |
| 23 | DIBER-23 | 2874 | LYG | Round | 7.27 | 5.12 | 6.8 | 51.32 | 17.6 | 31.08 | 25.5 | 12.1 | 2.1 | 342 | 2.17 | 3.01 |
|  | Range |  |  |  | **6.13-8.75** | **4.80-6.65** | **5.3-8.3** | **44.21-77.17** | **6.08-20.9** | **8.01-35.56** | **17.5-39.3** | **10-20.2** | **2.1-2.8** | **312-474** | **1.92-3.01** | **2.73-3.67** |
|  | Mean |  |  |  | 7.078 | 5.915 | 6.53 | 57.47 | 15.7 | 26.753 | 29.126 | 13.3 | 2.41 | 380.26 | 2.3491 | 3.11 |
|  | SEm± |  |  |  | 0.0721 | 0.0654 | 0.312 | 0.621 | 0.121 | 0.234 | 0.124 | 0.012 | 0.011 | 23.78 | 0.121 | 0.110 |
|  | CD (p=0.05) |  |  |  | 1.52 | 1.82 | 1.74 | 7.85 | 3.42 | 3.54 | 4.21 | 1.32 | 1.01 | 124.2 | 1.12 | 1.01 |

**Table-2.** Physico-chemical characteristics of fruits of *Hippophae salicifolia* morphotypes in Uttarakhand from Hanuman Chatti region

| S. No | Sample | Altitude (m) | Fruit colour And code | Fruit Shape | Fruit length (cm) | Fruit width (cm) | TSS (°B) | Juice (%) | Seed (%) | Pomace (%) | 100 fruit weight (g) | 100 fruit Volume (ml) | pH | Vitamin 'C' mg/100 g | Acidity % mg/100 g | Total Sugar mg/100g |
| --- | --- | --- | --- | --- | --- | --- | --- | --- | --- | --- | --- | --- | --- | --- | --- | --- |
| 1 | DIBER-24 | 2874 | LY | Round | 7.15 | 5.78 | 6.0 | 50.24 | 20.1 | 29.66 | 28.6 | 15.2 | 2.3 | 271 | 2.18 | 3.12 |
| 2 | DIBER-25 | 2874 | LY | Oval | 6.87 | 5.88 | 7.4 | 49.15 | 19.8 | 31.05 | 25.5 | 12.1 | 2.8 | 413 | 1.78 | 3.21 |
| 3 | DIBER-26 | 2552 | LG | Round | 6.56 | 6.66 | 7.2 | 61.19 | 16.9 | 21.91 | 34.7 | 16.2 | 2.7 | 342 | 2.37 | 3.11 |
| 4 | DIBER-27 | 2552 | LY | Round | 6.96 | 6.10 | 7.8 | 62.25 | 18.2 | 19.55 | 28.9 | 12.6 | 2.5 | 494 | 2.49 | 2.87 |
| 5 | DIBER-28 | 2552 | LY | Round | 7.11 | 6.21 | 7.0 | 56.62 | 19.6 | 23.78 | 33.9 | 15.2 | 2.6 | 376 | 1.86 | 2.98 |
| 6 | DIBER-29 | 2552 | LY | Round | 6.78 | 6.10 | 6.2 | 65.21 | 17.6 | 17.19 | 28.2 | 13.6 | 2.4 | 356 | 2.72 | 3.11 |
| 7 | DIBER-30 | 2552 | LY | Round | 6.14 | 5.89 | 6.4 | 64.14 | 18.4 | 17.46 | 25.6 | 11.8 | 2.3 | 294 | 2.08 | 3.23 |
|  | Range |  |  |  | 6.14-7.15 | 5.78-6.66 | 6.0-7.8 | 44.15-65.21 | 16.9-20.1 | 17.19-31.05 | 25.5-34.7 | 11.8-16.2 | 2.3-2.8 | 271-494 | 1.78-2.72 | 2.87-3.23 |
|  | Mean |  |  |  | 6.79571 | 6.088 | 6.857 | 58.4 | 18.65 | 22.942 | 29.342 | 13.814 | 2.51 | 363.71 | 2.2114 | 3.09 |
|  | SEm± |  |  |  | 0.0564 | 0.025 | 0.213 | 0.674 | 0.122 | 0.225 | 0.141 | 0.011 | 0.01 | 22.42 | 0.211 | 0.09 |
|  | CD (p=0.05) |  |  |  | 1.56 | 1.78 | 1.65 | 7.89 | 3.22 | 3.61 | 4.19 | 1.23 | 1.11 | 125.9 | 1.09 | 1.10 |

**Table 3:**
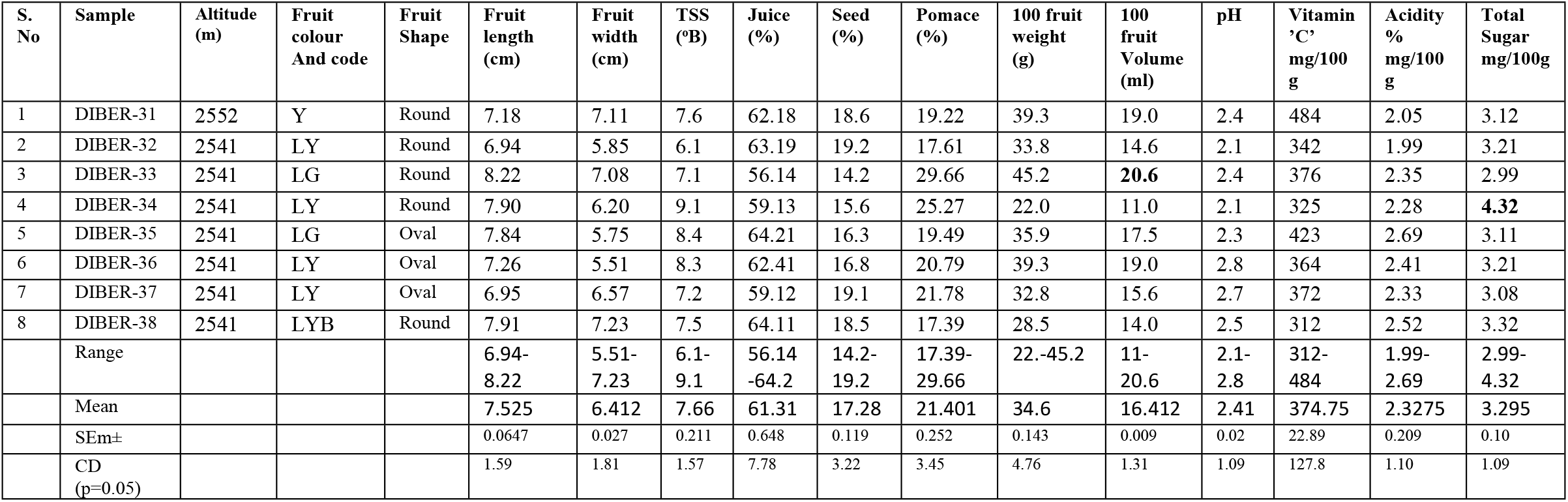
Physico-chemical characteristics of fruits of *Hippophae salicifolia* morphotypes from Auli region (Joshimath) of Uttarakhand

**Table 4:** Physico-chemical characteristics of fruits of *Hippophae salicifolia* morphotypes from Tharali region of Uttarakhand

| S.No | Sample | Altitude (m) | Fruit colour And code | Fruit Shape | Fruit length (cm) | Fruit width (cm) | TSS (°B) | Juice (%) | Seed (%) | Pomace (%) | 100 fruit weight (g) | 100 fruit Volume (ml) | pH | Vitamin 'C' mg/100 g | Acidity % mg/100 g | Total Sugar mg/100 g |
| --- | --- | --- | --- | --- | --- | --- | --- | --- | --- | --- | --- | --- | --- | --- | --- | --- |
| 1 | DIBER-39 | 2541 | LY | Round | 6.34 | 7.21 | 6.3 | 61.23 | 17.6 | 21.17 | **43.1** | 20.2 | 2.6 | 396 | 2.78 | 2.87 |
| 2 | DIBER-40 | 2541 | LY | Round | 8.74 | 7.59 | 8.4 | **76.19** | 15.5 | 8.31 | 17.5 | 10.0 | 2.4 | 474 | 1.98 | 2.99 |
| 3 | DIBER-41 | 2541 | LG | Round | 7.39 | 6.0 | 5.1 | 50.12 | 18.6 | 31.28 | 25.9 | 13.0 | 2.3 | 301 | 2.28 | 2.76 |
| 4 | DIBER-42 | 2541 | LY | Round | 7.30 | 5.48 | 9.1 | 59.54 | 17.5 | 22.96 | 28.8 | 13.5 | 2.4 | 376 | 2.21 | 2.87 |
| 5 | DIBER-43 | 2541 | LY | Oval | 8.14 | 6.46 | 7.9 | 61.12 | 18.2 | 20.68 | 27.2 | 12.8 | 2.1 | 403 | 1.92 | 3.01 |
| 6 | DIBER-44 | 2541 | LY | Oval | 7.3 | 5.8 | 8.4 | 62.14 | 19.4 | 18.46 | 32.4 | 14.2 | 2.3 | 356 | 2.05 | 3.23 |
| 7 | DIBER-45 | 2541 | LY | Round | 7.1 | 6.99 | 7.2 | 51.18 | 18.4 | 30.42 | 34.8 | 16.8 | 2.8 | 322 | 2.27 | 3.21 |
| 8 | DIBER-46 | 2541 | LY | Round | 7.87 | 7.12 | 7.5 | 50.29 | 16.8 | 32.91 | 24.5 | 12.0 | 2.7 | 437 | 1.95 | 2.98 |
| 9 | DIBER-47 | 2524 | LY | Round | 6.56 | 7.14 | 8.1 | 50.25 | 16.4 | 33.35 | 25.8 | 10.0 | 2.5 | 376 | 3.01 | 2.89 |
| 10 | DIBER-48 | 2524 | LY | Round | 8.77 | 7.42 | 7.1 | 51.27 | 17.9 | 30.83 | 27.3 | 13.0 | 2.2 | 305 | 2.15 | 3.04 |
| 11 | DIBER-49 | 2524 | LG | Round | 7.39 | 6.0 | 7.0 | 54.21 | 18.2 | 27.59 | 22.0 | 11.0 | 2.3 | 325 | 2.28 | 3.76 |
| 12 | DIBER-50 | 2524 | LY | Oval | 7.28 | 5.44 | 8.0 | 50.23 | 17.6 | 32.17 | 35.9 | 17.5 | **2.15** | 423 | 2.69 | 3.23 |
| 13 | DIBER-51 | 2524 | LG | Oval | 8.11 | 6.46 | 7.9 | 59.19 | 18.4 | 22.41 | 25.5 | 12.1 | 2.3 | 413 | 1.78 | 3.22 |
|  | Range |  |  |  | 6.34-8.74 | 5.44-7.59 | 5.1-9.0 | 50.12-76.19 | 15.5-19.4 | 8.31-33.35 | 17.5-43.1 | 10-20.2 | 2.1-2.8 | 305-474 | 1.78-3.01 | 2.76-3.76 |
|  | Mean |  |  |  | 7.5607 | 6.5469 | 7.53 | 56.689 | 17.7 | 25.58 | 28.51 | 13.54 | 2.38 | 377.4 | 2.2576 | 3.08 |
|  | SEm± |  |  |  | 0.0634 | 0.024 | 0.209 | 0.698 | 0.119 | 0.251 | 0.134 | 0.010 | 0.11 | 23.12 | 0.191 | 0.10 |
|  | CD (p=0.05) |  |  |  | 1.61 | 1.88 | 1.59 | 7.91 | 3.45 | 3.89 | 4.76 | 1.32 | 1.09 | 132.1 | 1.10 | 1.09 |

**Table 5:** Physico-chemical characteristics of fruits of *Hippophae salicifolia* morphotypes in Harshil region of Uttarakhand

| S. No | Sample | Altitude (m) | Fruit colour And code | Fruit Shape | Fruit length (cm) | Fruit width (cm) | TSS (°B) | Juice (%) | Seed (%) | Pomace (%) | 100 fruit weight (g) | 100 fruit Volume (ml) | pH | Vitamin' C' mg/100g | Acidity % mg/100g | Total Sugar mg/100g |
| --- | --- | --- | --- | --- | --- | --- | --- | --- | --- | --- | --- | --- | --- | --- | --- | --- |
| 1 | DIBER-52 | 2915 | LG | Oval | 6.47 | 6.08 | 6.9 | 61.18 | 18.6 | **38.30** | 34.7 | 16.2 | 2.2 | 342 | 2.37 | 3.04 |
| 2 | DIBER-53 | 2915 | LY | Round | 7.61 | 6.81 | 8.3 | 62.15 | 19.2 | 18.65 | 25.5 | 12.1 | 2.2 | 413 | 1.78 | 3.56 |
| 3 | DIBER-54 | 2915 | LY | Oval | 7.49 | 5.75 | 5.1 | 56.21 | 14.2 | 29.59 | 34.7 | 16.2 | 2.3 | 342 | 2.37 | 2.90 |
| 4 | DIBER-55 | 2915 | LYB | Oval | 7.41 | 4.93 | 9.6 | 65.17 | 15.6 | 19.23 | 28.9 | 12.6 | 2.4 | 494 | 2.49 | 2.89 |
| 5 | DIBER-56 | 2915 | LG | Round | 7.37 | 4.85 | 7.6 | 51.12 | 18.4 | 30.48 | 33.9 | 15.2 | 2.45 | 376 | 1.86 | 2.98 |
| 6 | DIBER-57 | 2915 | LY | Round | 7.51 | 6.42 | 8.7 | 50.21 | 16.8 | 33.08 | 28.2 | 13.6 | 2.1 | 356 | 2.72 | 3.09 |
| 7 | DIBER-58 | 2915 | LG | Round | 5.61 | 4.81 | 9.7 | 50.54 | 16.4 | 33.06 | 25.6 | 11.8 | 2.3 | 294 | 2.08 | 3.03 |
| 8 | DIBER-59 | 2786 | LY | Round | 6.45 | 5.89 | 8.1 | 51.74 | 17.9 | 30.36 | 24.5 | 12.0 | 2.8 | 437 | 1.95 | 3.09 |
| 9 | DIBER-60 | 2786 | LY | Oval | 6.45 | 5.89 | 8.1 | 54.47 | 18.2 | 27.33 | 24.5 | 12.0 | 2.7 | 437 | 1.95 | 2.98 |
| 10 | DIBER-61 | 2786 | LY | Oval | 6.14 | 5.40 | 7.6 | 51.52 | 18.4 | 30.08 | 25.8 | 10.0 | 2.5 | 376 | 3.01 | 2.87 |
|  | Range |  |  |  | 5.61-7.61 | 4.93-6.81 | 5.1-9.7 | 50.21-65.17 | 14.2-19.2 | 18.65-38.30 | 24.5-34.7 | 10-16.2 | 2.1-2.8 | 294-494 | 1.78-3.01 | 2.87-3.56 |
|  | Mean |  |  |  | 6.851 | 5.683 | 7.97 | 55.43 | 17.3 | 29.01 | 28.6 | 13.17 | 2.39 | 386.7 | 2.25 | 3.043 |
|  | SEm± |  |  |  | 0.0614 | 0.031 | 0.209 | 0.714 | 0.202 | 0.234 | 0.151 | 0.021 | 0.02 | 23.26 | 0.197 | 0.10 |
|  | CD (p=0.05) |  |  |  | 1.62 | 1.83 | 1.59 | 7.91 | 3.67 | 3.98 | 4.76 | 1.43 | 1.09 | 134.8 | 1.10 | 1.00 |

**Table 6:**
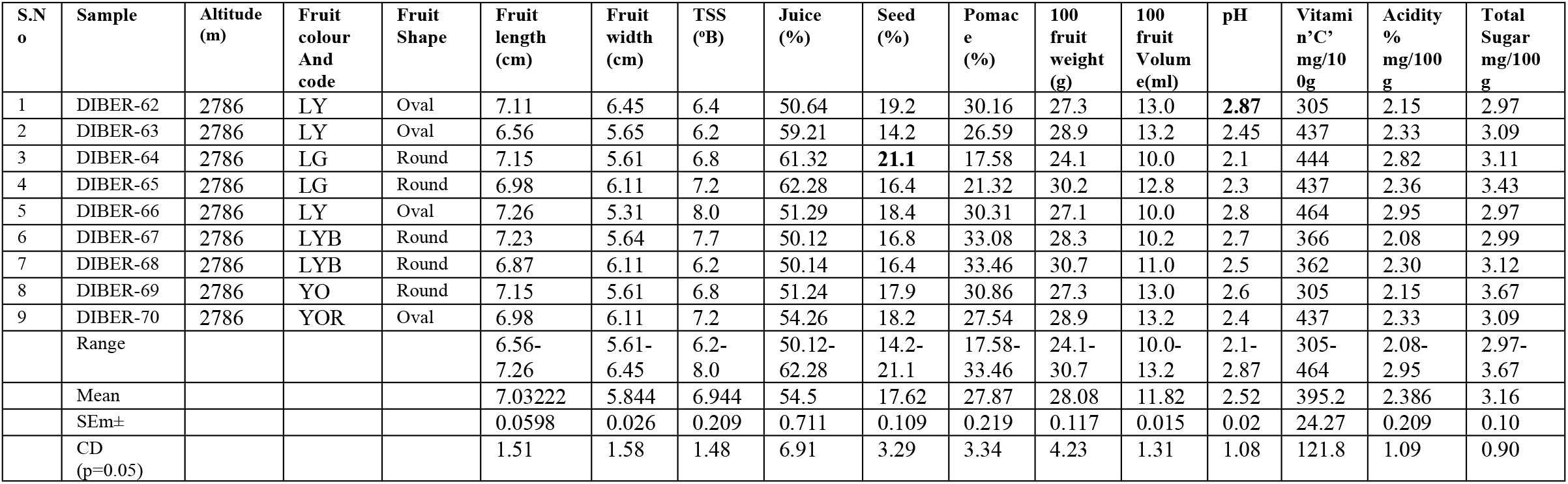
Physico-chemical characteristics of fruits of *Hippophae salicifolia* morphotypes from Urgam Valley in Uttarakhand

**Figure 1:**
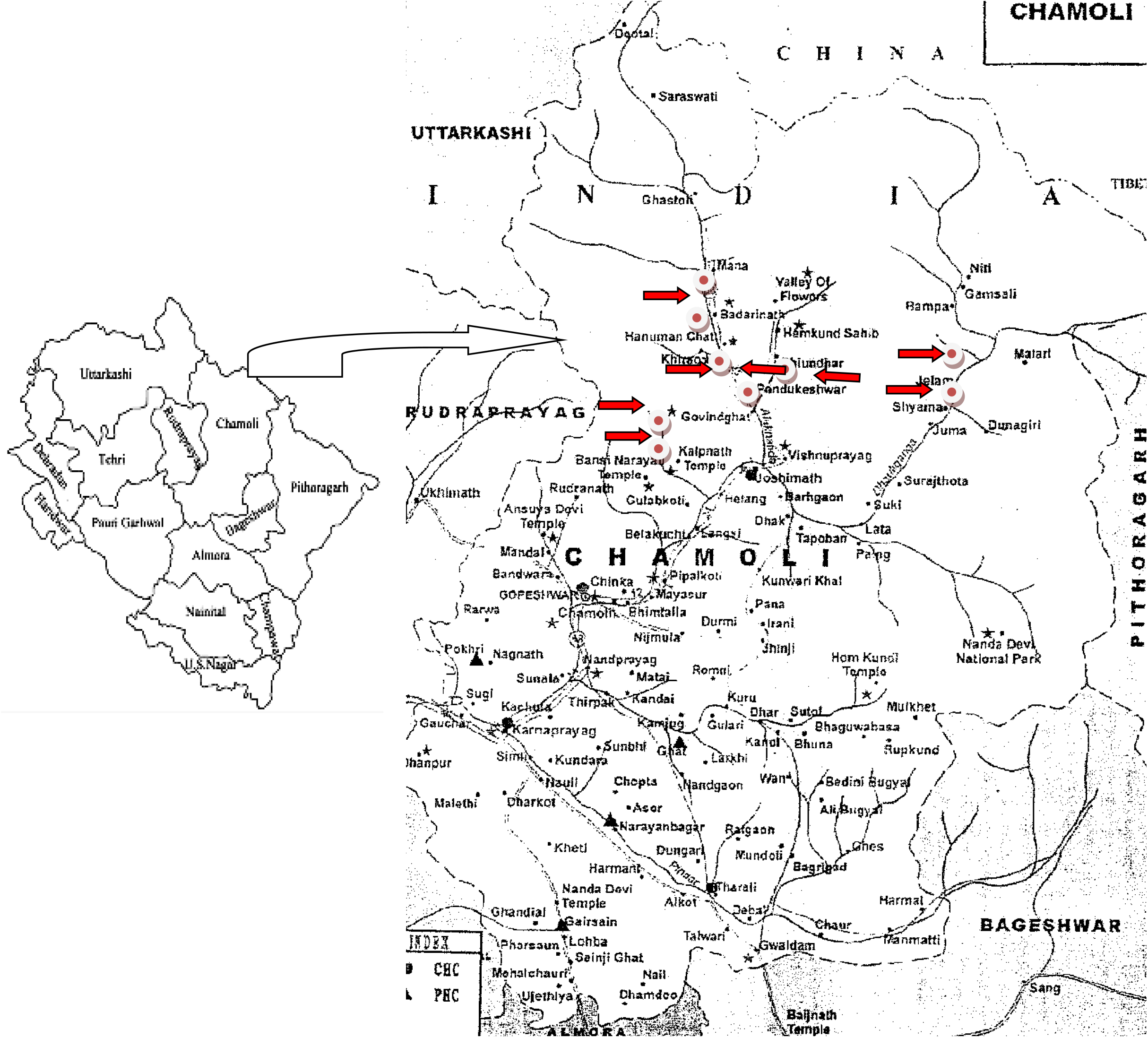
Geographical locations of Seabuckthorn Fruit collection sites in Chamoli District of Uttarakhand (India)

### Variation in Physical characteristics of fruits

Variation in the fruit parameters in the natural plant population of Seabuckthorn will be highly helpful in the selective evaluation of the plants with desirable traits, important for pharmaceutical and nutraceutical importance.Variability studies in the shape, size, color, weight and volume of the fruits and leaves were carried out and lot of variations were observed (Figure 2). Color of the fruit samples were recorded with the help of The Royal Horticultural society color chart and color varied from light yellow, yellow, yellow orange to orange and orange red in color. Shape of the fruit varied from oval, round, elliptical to capsular. Size of fruits of seabuckthorn is the limiting factor for its commercialization because of its small size and high perish-ability. It is obvious from data that fruit length was in the range of 6.14 (DIBER-30) to 8.77 mm (DIBER-48), with a mean value of 7.45 mm while fruit width ranged from 4.80mm (DIBER-8) to 7.59 mm (DIBER-40) with a mean of 6.19 mm. Maximum weight of 100 fruits was recorded 45.2g in (DIBER-33) whereas minimum was 17.5 g in (DIBER-8) with a mean value of 31.35 g. Similarly volume of 100 fruits showed variation from 10ml in ‘DIBER-8’ to 20.6 ml in ‘DIBER-33’ with a mean volume of 15.3 ml. Size, colour, appearance and weight of fruits are important characteristics of commercial cultivation and processing of *Hippophae salicifolia*. Plants having attractive fruit color and large size fruits have been selected since size and weight of the fruits are yield attributing characteristics (Nath et al. 2000). Therefore promising selections have been made considering these attributes.

**Figure 2.**
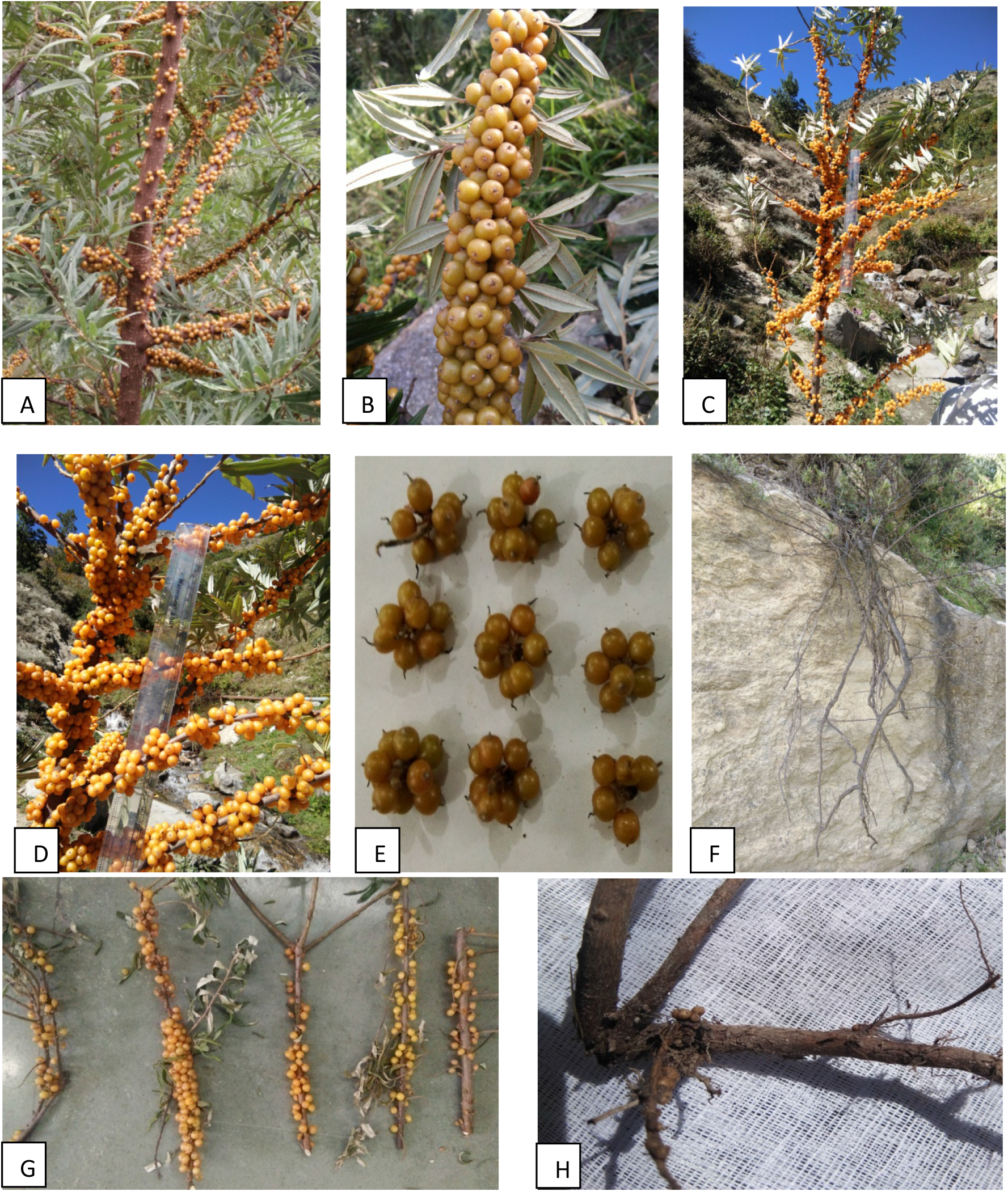
A-D. Fruit bearing branches of Seabuckthorn, E-Bunching pattern of fruits, F-Root length of underground root of seabuckthorn, G-Branches bearing different types of fruits, H-Root bearing root nodules.

**Figure 3.**
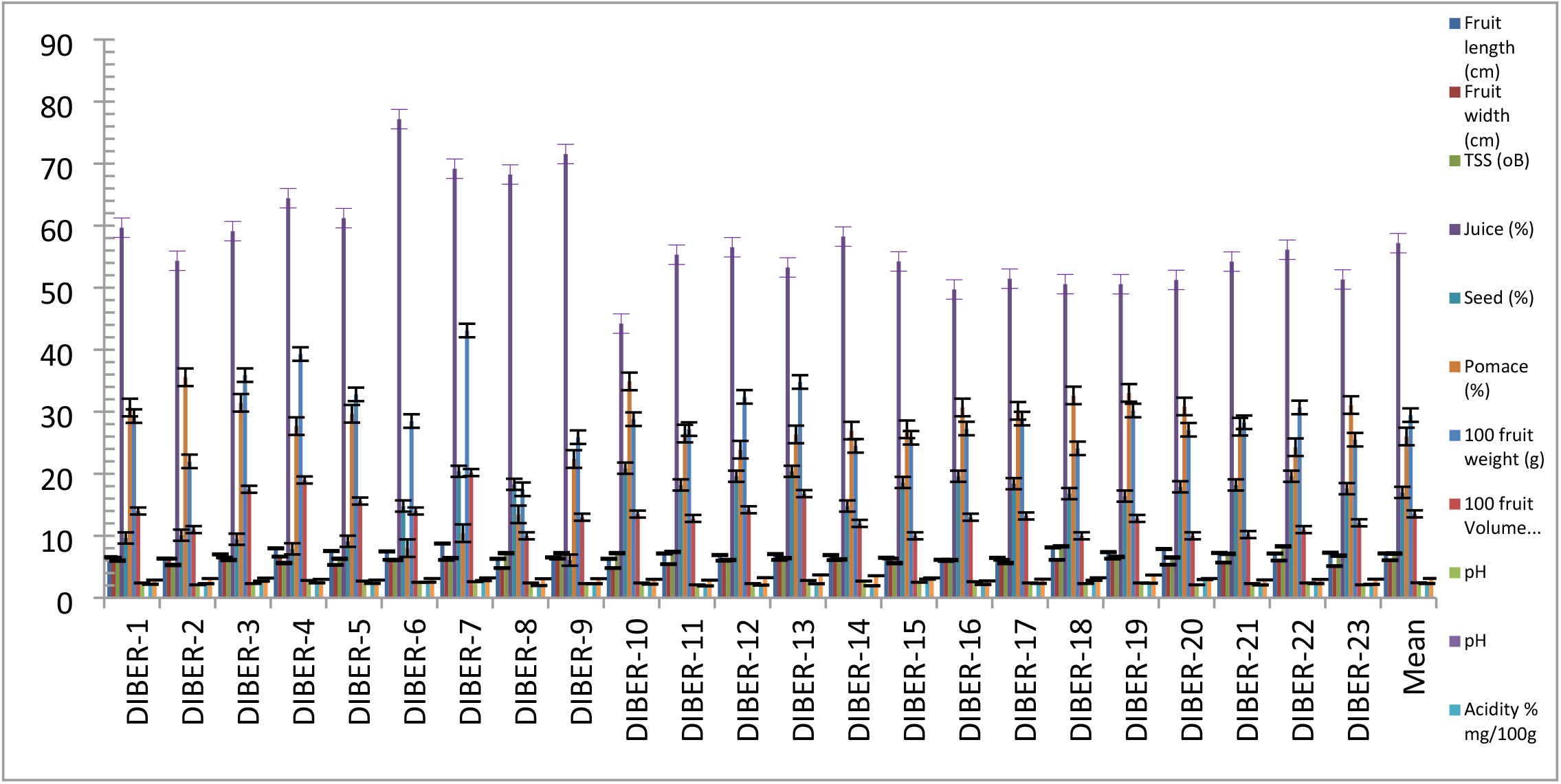
Physico-chemical characteristics of fruits of *Hippophae salicifolia* morphotypes (DIBER 1-23) of Uttarakhand

**Figure 4.**
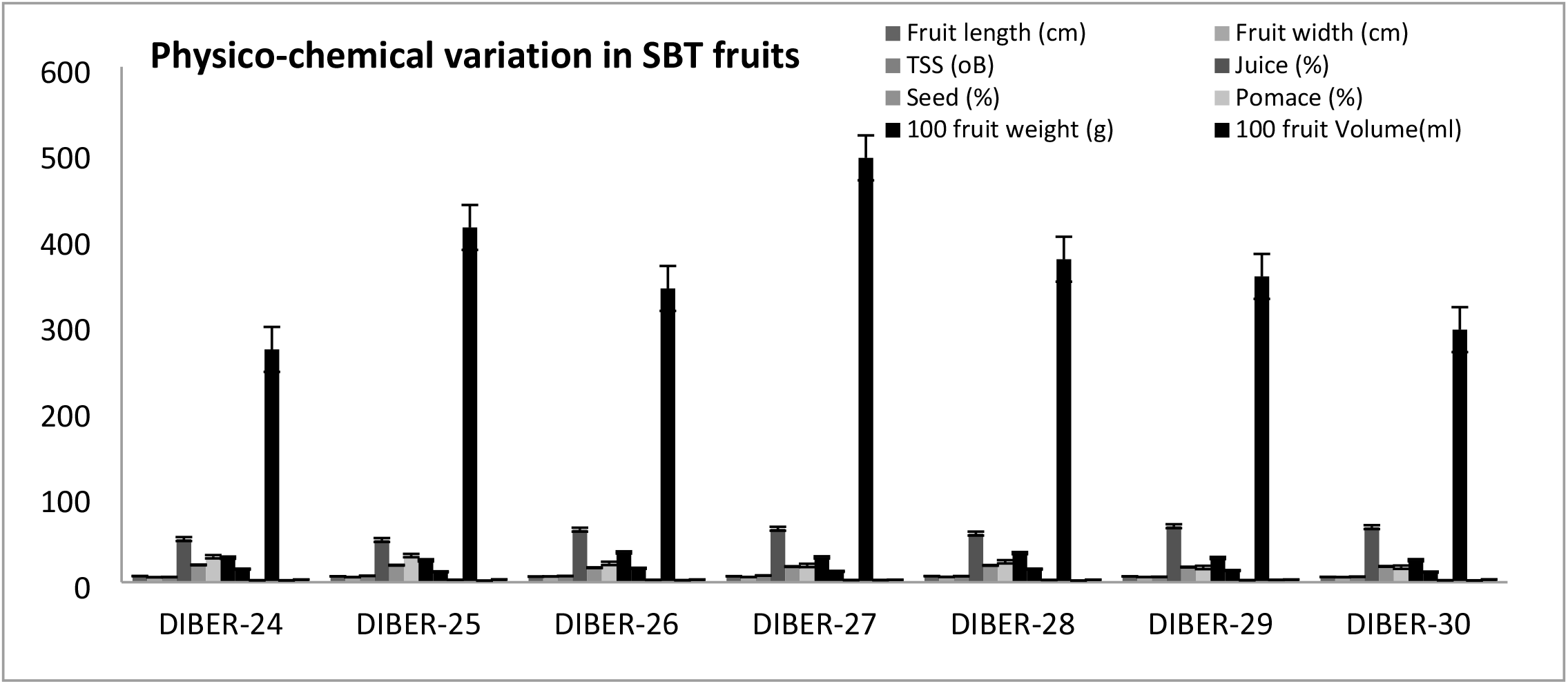
Physico-chemical characteristics of fruits of *Hippophae salicifolia* morphotypes (DIBER 24-30) of Uttarakhand

**Figure 4.**
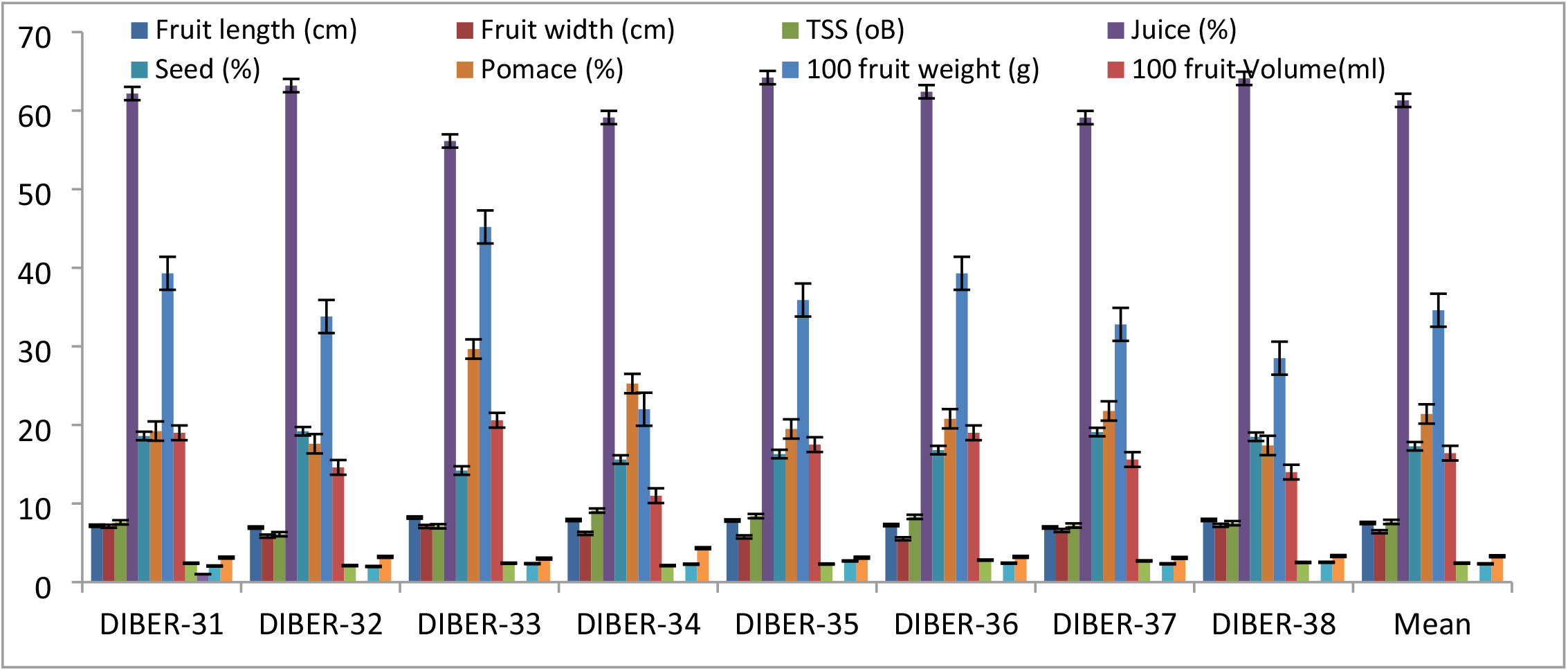
Physico-chemical characteristics of fruits of *Hippophae salicifolia* morphotypes (DIBER 31-38) of Uttarakhand

**Figure 4.**
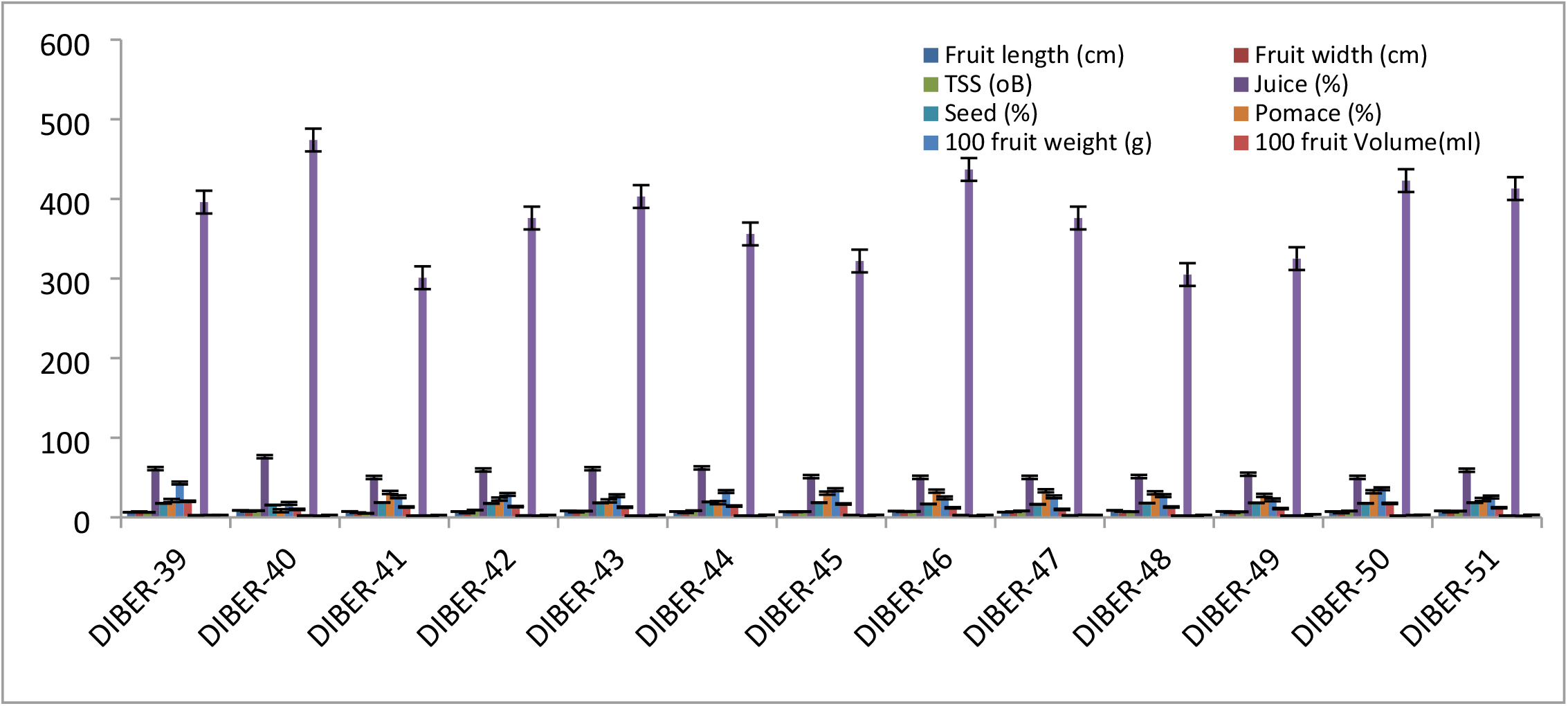
Physico-chemical characteristics of fruits of *Hippophae salicifolia* morphotypes (DIBER 39-51) of Uttarakhand

**Figure 5.**
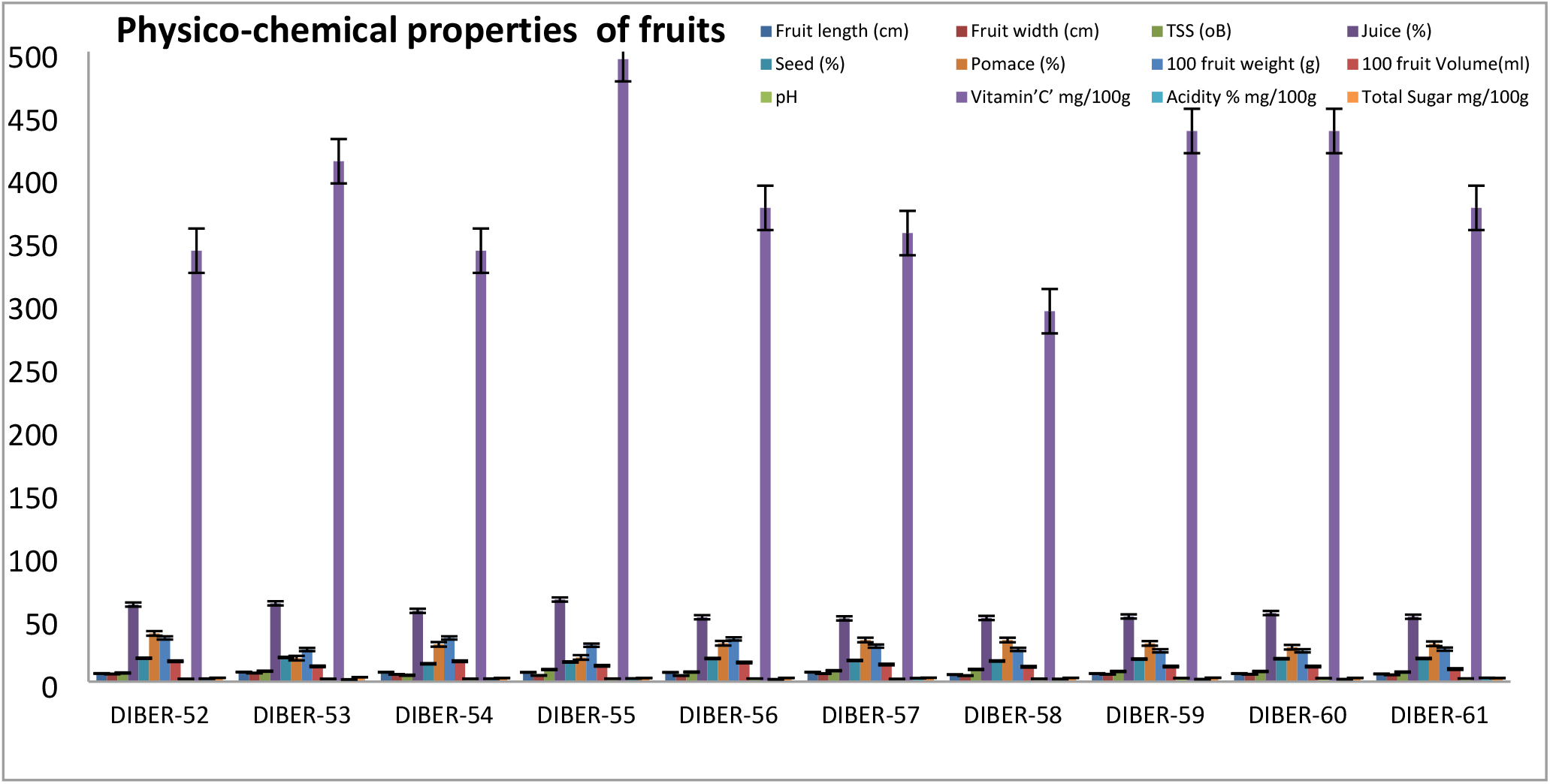
Physico-chemical characteristics of fruits of *Hippophae salicifolia* morphotypes (DIBER 52-61) of Uttarakhand

**Figure 6.**
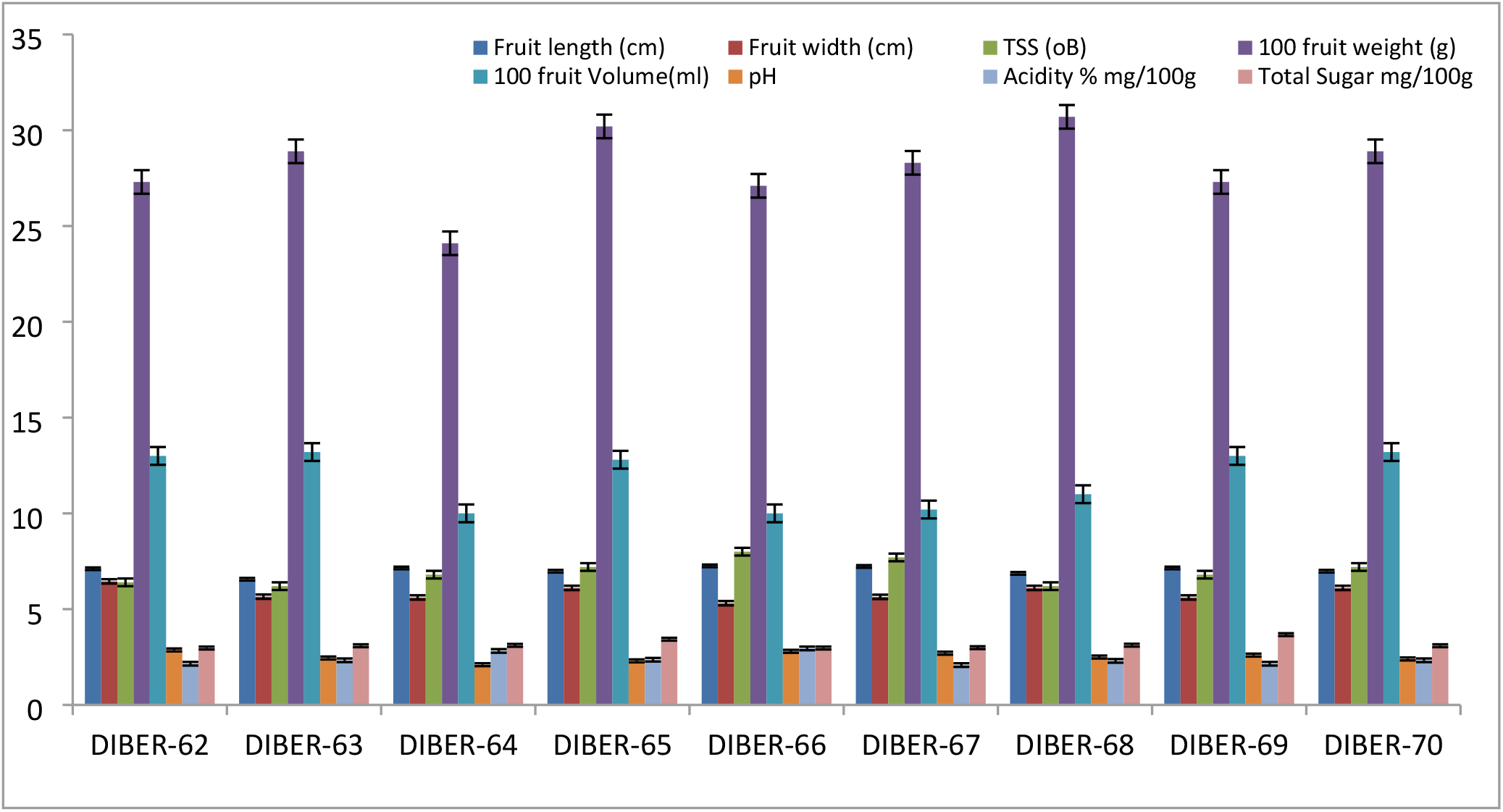
Physico-chemical characteristics of fruits of *Hippophae salicifolia* morphotypes (DIBER 62-70) of Uttarakhand

## Variation in chemical properties of fruits

Physico-chemical properties like pH, TSS, acidity, sugars and vitamin ‘C’ content of *Hippophae salicifolia* are essential parameters for nutritional and processing purposes. All these parameters have shown variation in the fruit samples (Table-1) The total soluble solids (TSS) varied from 5.1in ‘DIBER-41’ to 9.7 in ‘DIBER-58’ with a mean value of 7.4oB and pH of juice ranged from 2.15 to 2.87 in ‘Diber-50’ and ‘Diber-62’ respectively with a mean value of 2.51. Acidity of the fruit juice was in the range of 1.92 to 3.58 % in DIBER-51 and DIBER-69 respectively with a mean value of 2.75 %. Vitamin ‘C’ was found highest 494mg/100g in DIBER-55 and lowest 271 mg/100g in DIBER-24 with a mean value of 382.5 mg/100g of fruits. Total sugar content in fruit pulp was minimum 2.73% in DIBER-16 and highest 4.32% in DIBER-34 with a mean value of 3.53%. Based on these parameters the promising genotypes having highest vitamin ‘C’ and TSS have been selected for multiplication and conservation.

### Variation in Juice, seed and pomace percentage of fruits

*Hippophae salicifolia* fruits being highly acidic, soft and perishable in nature are not consumed fresh for table purposes in general. Processing of fresh fruits soon after harvesting thus becomes imperative. For selection of improved genotypes of *Hippophae salicifolia*, data on Juice, Seed and Pomace content in fruit are important for establishing their suitability for processing. Among the various fruit samples collected Fruit Juice % varied from 44.21% in DIBER-10 to 76.19 %, in DIBER-40 with a mean value of 60.20%. Pomace ranges from 8.01% in DIBER-6 to 28.30 % in DIBER-51 with a mean value of 18.15 % and seed % was in the range of 6.8% in DIBER-9 to 21.1 % in DIBER-65 with a mean value of 13.95%. It is obvious from the data that there is distinct variation in the juice content of the fruit which might have been influenced by growing place and genetic makeup of the morphotypes (Dwivedi et al, 2004). Seed and pomace are the major residues after processing of *Hippophae salicifolia* fruits which contains vitamins and valuable oil suitable for medicinal and cosmetic industries (Rongsen, 1992).

Among the 70 fruit samples collected from Chamoli District of Uttarakhand, 6 promising genotypes have been identified based on desirable characteristics. These selected genotypes have been conserved at DIBER field station, Auli (3142m above mean sea level) which can be utilized in *Hippophae salicifolia* breeding programme as parent material.

## Acknowledgement

Authors are highly thankful to Dr. Basant Ballabh, TO ‘B’ and Sh. H. B. Mehra, STA ‘B’ for their valuable support during the survey and sample collection of seabuckthorn growing areas.

